# PTTG1-Mediated Pericyte Dysfunction Drives Diabetes-Induced Microvascular Dysfunction

**DOI:** 10.1101/2024.12.02.626502

**Authors:** Linyu Zhang, Ling Ren, Jingyue Zhang, Min Xia, Xiaosa Li, Mudi Yao, Fanfei Ma, Chang Jiang, Jin Yao, Biao Yan

## Abstract

**Background:** Pericytes are crucial for the development, stabilization, and functional regulation of microvasculature, especially in the retina. In diabetic retinopathy (DR), early loss of pericytes is a key event that drives microvascular dysfunction. Despite their critical role, the mechanisms underlying the functional heterogeneity of pericytes in DR remain poorly understood, impeding the development of effective therapeutic strategies.

**Methods:** We employed single-cell RNA sequencing to construct a comprehensive single-cell atlas of non-diabetic and diabetic retinas. Using bioinformatic clustering and subcluster analysis, we identified a specific pericyte subcluster associated with diabetic microvascular complications. Differential gene expression analysis and immunofluorescence validation highlighted PTTG1 as a potential key regulator of pericyte dysfunction. To investigate its functional role, we emplyed CRISPR/Cas9 and adenoviral vectors to modulate PTTG1 expression in vitro and in vivo. Combined transcriptomic and metabolomic approaches were used to explore the mechanistic pathways through which PTTG1 influences pericyte biology and vascular function.

**Results:** We identified a novel pericyte subcluster characterized by elevated expression of PTTG1, which was strongly correlated with diabetic microvascular dysfunction. Silencing PTTG1 using CRISPR/Cas9 and siRNA in vitro mitigated pericyte dysfunction under high-glucose conditions. Targeted knockdown of PTTG1 using viral vectors improved retinal vascular integrity and reduced neovascularization in diabetic mice. Transcriptomic and untargeted metabolomic analyses revealed that PTTG1 knockdown reprogrammed pericyte energy metabolism by modulating glycolysis pathway genes, reducing oxidative stress, and restoring pericyte function, ultimately alleviating microvascular dysfunction in DR.

**Conclusions:** PTTG1 plays a critical role in regulating pericyte dysfunction and maintaining vascular homeostasis in diabetic retinopathy. By modulating key metabolic pathways and pericyte phenotypes, PTTG1 represents a promising therapeutic target for treating diabetic microvascular complications. These insights offer a novel molecular framework for developing targeted therapies aimed at restoring retinal vascular health in diabetic patients.

## Introduction

Diabetes mellitus, a widespread chronic metabolic disorder, has emerged as a major global public health concern due to its high prevalence and association with multiple life-threatening complications.^1^ Persistent hyperglycemia and genetic predisposition lead to systemic microvascular dysfunction, ultimately resulting in severe complications such as blindness, end-stage renal disease, cerebrovascular events, cardiomyopathy, and peripheral neuropathy with necrosis.^2,3^ Among them, diabetic retinopathy (DR) is the most common and widespread microvascular complication of diabetes, serving as a key model for studying diabetes-related microvascular dysfunction.^4^ In DR, microarterial damage is initially characterized by microaneurysm formation, increased vascular permeability, and cotton wool spots. In advanced stages, pathological retinal neovascularization and fibrovascular membrane proliferation lead to severe complications, such as retinal detachment and subsequent vision loss.^5^ Although microvascular injury affects various organs differently, it shares a key early hallmark: pericyte apoptosis and loss.^6^ Pericytes are a unique type of mural cell that is present in the vasculature throughout the body. They are embedded within the same basement membrane as endothelial cells (ECs) and regulate normal vascular functions.^7,8^ The intimate association between pericytes and endothelial cells results in progressive endothelial cell loss, dysregulation of vascular tone, and ischemic alterations.^9–11^ Therefore, understanding the changes in pericytes and vasculature in DR offers a valuable window into the systemic vascular pathology of diabetes, providing a scientific basis for developing early intervention and treatment strategies for diabetic microvascular complications. Previous studies found that factors inducing pericyte apoptosis in a hyperglycemic environment included increased production of reactive oxygen species, accumulation of advanced glycation end products, and downregulation of the expression or activity of antioxidant proteins.^12,13^ However, to date, research on how pericytes mobilize signaling pathways and undergo metabolic shifts in an oxidative stress environment of diabetes mellitus remains significantly underexplored compared to other vascular cells.

In recent years, there has been a growing recognition of the substantial functional heterogeneity exhibited by pericytes. In the glomerulus, pericytes function as mesangial cells that closely encase the microvascular walls, regulating filtration flow.^14^ In the central nervous system, pericytes are critical components of the blood-brain barrier (BBB), while in skeletal muscle, they regulate myogenesis and can differentiate into fibroblasts.^15,16^ In the cardiovascular and respiratory systems, pericytes play vital roles in regulating angiogenesis and facilitating vascular repair.^17,18^ Notably, retinal vasculature has a high pericyte coverage. Pericytes with different functions exhibit distinct gene expression profiles and morphologies, contributing to angiogenesis, vascular stabilization, the formation of the blood-retinal barrier, blood flow regulation, and neurovascular coupling.^19^ This functional heterogeneity of pericytes offers new perspectives for in-depth research into the mechanisms of systemic vascular pathology in diabetes. However, studies on the definition and function of heterogeneous pericyte subclusters remain in the early stages.

Investigating pericyte heterogeneity offers new avenues for developing therapies that target specific pericyte subtypes associated with various diseases. By intervening in functional subgroups of pericytes, it may become feasible to precisely address the onset and progression of DR and other diabetic microvascular complications. Single-cell RNA sequencing allows the study of regulatory mechanisms of relevant cell subtypes and states in DR based on the specific gene profiles of individual cells. By analyzing differential expression in various pericyte populations with similar transcriptomes, key genes associated with each cell type or state can be identified.^20–22^ Single-cell sequencing enables the discovery of functional characteristics and gene expression differences among pericyte subpopulations involved in diabetic vascular dysfunction.

In this study, we integrated single-cell RNA sequencing analysis with *in vitro* and *in vivo* phenotypic studies to investigate pericyte heterogeneity in the context of DR and to explore potential therapeutic strategies targeting pericyte dysfunction. We identified three pericyte subclusters, with subcluster 0 closely associated with pathological angiogenesis. Notably, this subcluster was characterized by high expression of pituitary tumor transforming gene 1 (PTTG1), which was also significantly upregulated in the DR group. Subsequent experiments confirmed that PTTG1 expression was significantly increased in pericytes under diabetic stress, and silencing PTTG1 could alleviate retinal vascular dysfunction. Mechanistic studies revealed that a shift in glycolytic metabolism influenced the apoptotic process of pericytes. This study offers new insights into the transcriptomic heterogeneity of retinal microvascular pericytes. Therapeutic interventions targeting pericyte PTTG1 regulation may provide a new approach to prevent and protect against diabetes-induced retinal vascular damage.

## Methods

### Data Availability

All supporting data are provided within the article and the Supplemental Material.

## Results

### Generation of the single-cell atlas of non-diabetic and diabetic retinas

To investigate pericyte heterogeneity during the pathological processes of DR, we established a DR mouse model by intraperitoneal injection of streptozotocin (STZ). Once the model was confirmed, both non-diabetic and diabetic mice were euthanized, and their retinas were isolated to prepare single-cell suspensions. Subsequently, the samples underwent single-cell RNA sequencing, followed by comprehensive data analysis. The whole process is illustrated in Figure 1A. After obtaining the data, we conducted quality filtering and batch correction as initial steps. The criteria for filtering, including gene number in each cell (nFeature_RNA), total count of all genes in each cell (nCount_RNA), and percentage of mitochondrial genes in total genes in each cell (percent.mt), were established and presented in Figure 1B. The correlation coefficient between nFeature_RNA and nCount_RNA was found to be 0.91 in both non-diabetic and diabetic groups (Figure 1C).

**Figure 1.**
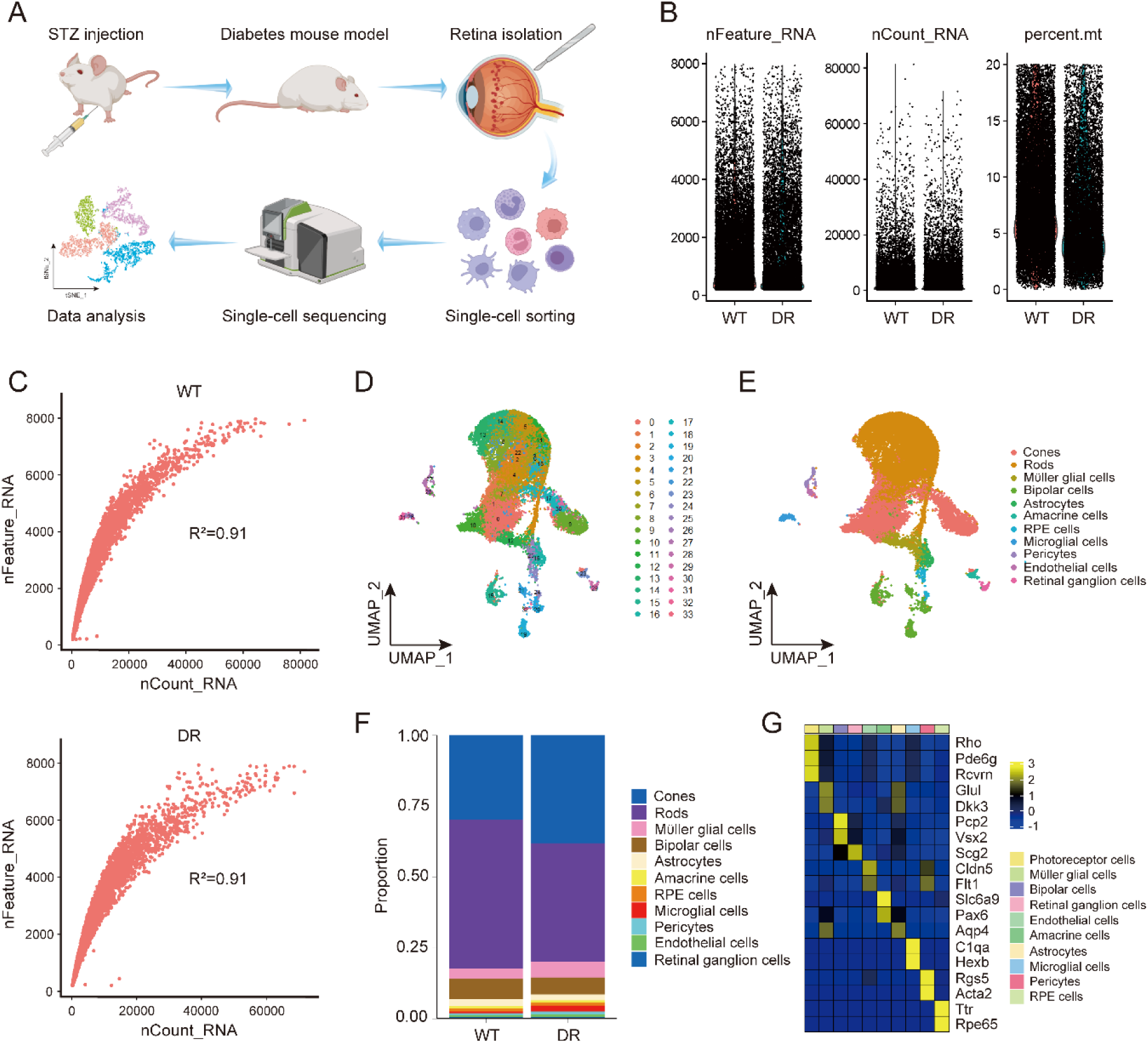
A single-cell atlas of non-diabetic and diabetic retinas. (A) Illustration shows the process of generating a single-cell transcriptome atlas. The DR model was induced by STZ injection, and after 25 weeks, retinas were isolated to prepare single-cell suspensions. Libraries were sequenced, and data analyzed. (B) Quality control criteria for each cell are shown, including gene count per cell (nFeature_RNA), total gene count (nCount_RNA), and mitochondrial gene percentage (percent.mt). (C) Seurat’s Pearson correlation analysis reveals relationships between nFeature_RNA and nCount_RNA. (D, E) UMAP plots display 34 cell clusters and identified cell types, such as photoreceptors, Müller glia, RPE cells, microglia, pericytes, ECs, and ganglion cells. (F) Barplot compares retinal cell type proportions in non-diabetic (WT) and diabetic (DR) samples. (G) Heatmap shows average expression of cell-type-specific markers validating clusters.

Following quality control filtering, we obtained 18,926 cells from the non-diabetic group and 19,056 cells from the diabetic group, respectively. Initially, these cells were classified into 34 transcriptionally distinct clusters and visualized in a two-dimensional space using the UMAP algorithm (Figure 1D). To annotate cell types, we analyzed the average expression levels of established cellular markers in each cluster. Subsequently, we identified 10 cell types (Figure 1E), including photoreceptor cells (Rho, Pde6g, Rcvrn), Müller glial cells (Glul, Dkk3), bipolar cells (Pcp2, Vsx2), astrocytes (Aqp4), amacrine cells (Slc6a9, Pax6), retinal pigment epithelial (RPE) cells (Ttr, Rpe65), microglial cells (C1qa, Hexb), pericytes (Rgs5, Acta2), ECs (Cldn5, Flt1), and retinal ganglion cells (Scg2). The average expression levels of selected cell-type-specific markers used for validating each cell cluster were depicted in Figure 1G. Additionally, we calculated and compared the proportions of each cell type between the non-diabetic and diabetic groups (Figure 1F).

### Pericyte subcluster 0 is potentially associated with pathological angiogenesis in diabetic retina

To enhance our comprehension of the transcriptomic heterogeneity of pericytes during the pathological angiogenesis in DR, we subsequently focused on pericytes and performed UMAP reduction analysis on this cell type. Three distinct pericyte subclusters were identified, and the proportions of cells from non-diabetic and diabetic groups in each subcluster were compared (Figure 2A and Figure S1A). The UMAP plot reveals distinct transcriptional profiles for pericyte subclusters 0, 1, and 2, suggesting potential differences in their biological functions. The highly expressed genes in each subcluster are depicted in Figure 2B and Figure S1B.

**Figure 2.**
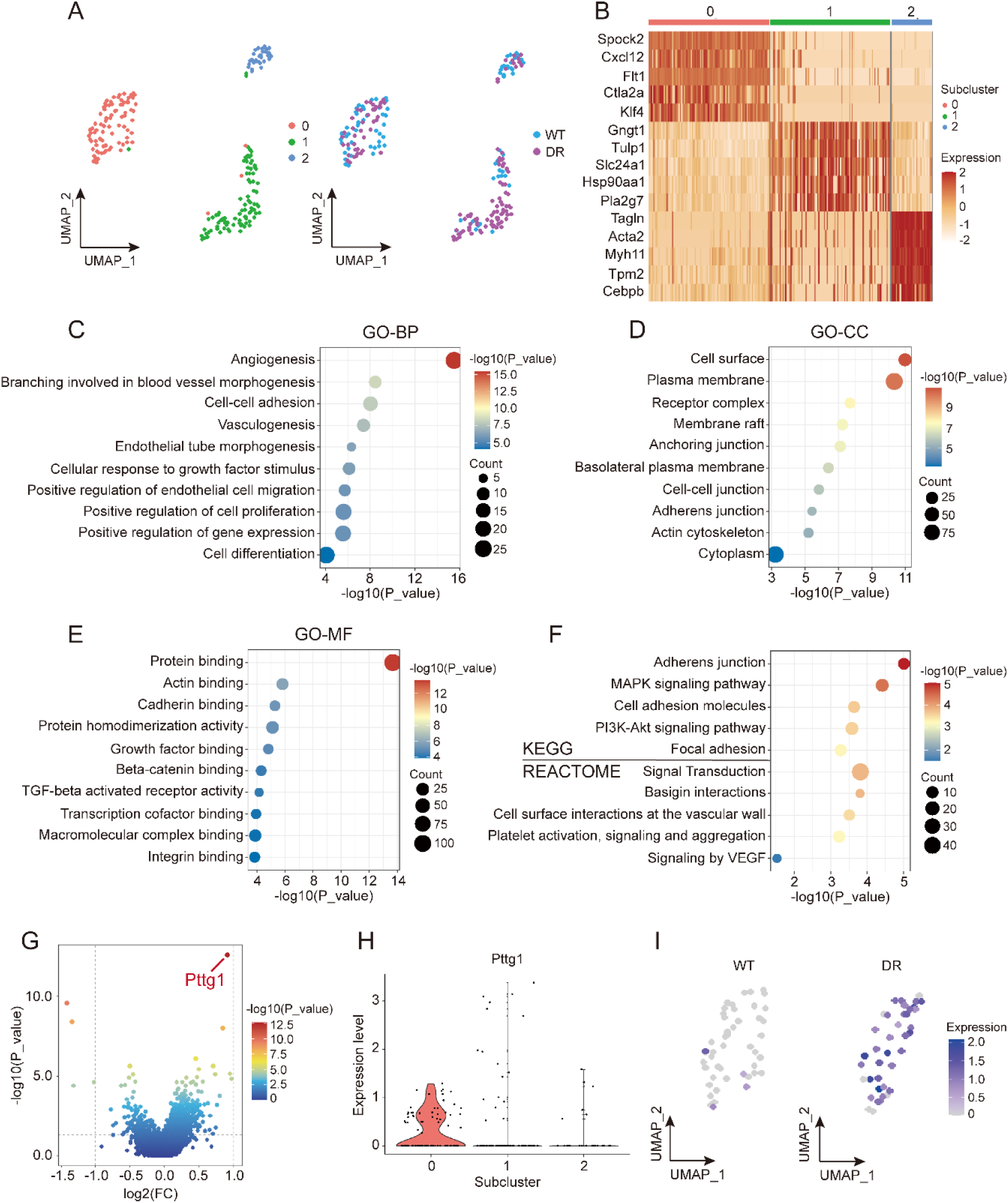
The unique functions and differentially expressed genes of pericyte subcluster 0 (A) UMAP plots show three distinct pericyte subclusters in non-diabetic and diabetic retinas. (B) Heatmap depicts the top 5 highly expressed genes in each pericyte subcluster. (C-E) Dot plots represent GO enrichment of the top 200 highly expressed genes in pericyte subcluster 0, categorized into biological process (C), cellular component (D), and molecular function (E). (F) Dot plot presents pathway enrichment results from KEGG and REACTOME databases using the top 200 highly expressed genes in subcluster 0. (G) Volcano plot highlights differentially expressed genes between diabetic and non-diabetic groups in subcluster 0. (H) Violin plot illustrates PTTG1 expression across pericyte subclusters. (I) FeaturePlots show PTTG1 expression in diabetic and non-diabetic groups within subcluster 0.

Subcluster 0 exhibited a high abundance of genes associated with cell adhesion, cell proliferation, extracellular matrix binding, VEGF signaling, and regulation of angiogenesis, including Spock2, Cxcl12, Flt1, and Klf4, indicating that subcluster 0 is actively involved in EC-pericyte interaction and pathological angiogenesis. In subcluster 1, Tulp1, Hsp90aa1, and Pla2g7 are known to be involved in stimulation of phagocytosis, cell response to environmental stress, and regulation of inflammatory and oxidative stress response, implying that subcluster 1 is potentially associated with vascular inflammatory processes in DR. Subcluster 2 highly expressed Tagln, Acta2, and Tpm2, which are essential for smooth muscle differentiation and contraction, vascular contractility, cell motility and actin binding, suggesting that pericytes in subcluster 2 are responsible for the maintenance of normal vascular function. Based on the results, we can infer that pericyte subcluster 0 is potentially associated with pathological angiogenesis in DR.

### The unique functions and differentially expressed genes of pericyte subcluster 0

To elucidate the potential functions of pericyte subcluster 0, we conducted Gene Ontology (GO) enrichment analysis using the top 200 highly expressed genes within the subcluster, encompassing GO-Biological Process (GO-BP), GO-Cellular Component (GO- CC), and GO-Molecular Function (GO-MF) terms (Figure 2C - 2E). GO-BP analysis revealed enrichments tightly associated with neovascularization processes, including ’Angiogenesis,’ ’Branching involved in blood vessel morphogenesis,’ ’Cell-cell adhesion,’ ’Vasculogenesis,’ ’Endothelial tube morphogenesis,’ ’Cellular response to growth factor stimulus,’ ’Positive regulation of EC migration,’ and ’Positive regulation of cell proliferation.’ GO-CC results showed that the top 200 highly expressed genes in subcluster 0 were mainly enriched in locations of cell membrane, cell junction, and cytoplasm. GO- MF analysis also revealed terms linked to cell proliferation and cell adhesion, including ’Actin binding,’ ’Cadherin binding,’ ’Growth factor binding,’ ’Beta-catenin binding,’ ’TGF-beta activated receptor activity,’ and ’Integrin binding.’ Moreover, pathway enrichment analysis using KEGG and REACTOME databases suggested that these highly expressed genes were significantly enriched in pathways related to cell adhesion, the MAPK signaling pathway, the PI3K-Akt signaling pathway, and cell surface interactions (Figure 2F). These findings further support the close association between pericyte subcluster 0 and pathological angiogenesis.

To investigate the transcriptomic changes in the diabetic group within subcluster 0, we compared the expression profiles between non-diabetic and diabetic groups and utilized a Volcano plot to visualize the differentially expressed genes (Figure 2G). PTTG1 emerged as the most significantly upregulated gene in the diabetic group. Notably, Violin plot showed that PTTG1 is dominantly expressed in pericyte subcluster 0 (Figure 2H), and FeaturePlots further evidenced the significant upregulation of PTTG1 in the diabetic group (Figure 2I). These results suggest a potential association between PTTG1 and pathological angiogenesis of DR.

### PTTG1 expression is increased in diabetic retina and highly expressed in pericytes

Since PTTG1 was found to be upregulated in the DR group of pericyte subcluster 0, the retinas from non-diabetic and STZ-induced DR mice were sampled to detect PTTG1 expression. qRT-PCR and western blotting analyses revealed a considerably elevated PTTG1 expression in the retinas of DR mice (Figure 3A, B). Compared with non-diabetic mice, DR mice showed significantly increased PTTG1 expression in retinal tissues, which co-localized with the retinal microvascular pericyte markers, chondroitin sulfate proteoglycan (NG2) (Figure 3C). Furthermore, in retinal cryosections, immunofluorescence assays showed co-localization of PTTG1 with the pericyte marker NG2, which was consistent with the bioinformatics analysis (Figure 3D, E). To further ascertain the location of PTTG1 expression, we also confirmed that PTTG1 was not co-localized with ganglion cell markers RNA Binding Protein with Multiple Splicing (RBPMS), nor with glial cell marker Glutamine Synthetase (GS) (Figure S2A-D). To further validate the association between PTTG1 and pathological angiogenesis in retina, we also performed immunofluorescence staining in oxygen-induced retinopathy (OIR) mouse model. The results showed that PTTG1 expression was increased in pathological angiogenic regions of retinas (Figure S2E). These results suggest that PTTG1 may participate in pathological angiogenesis through regulating pericytes.

**Figure 3:**
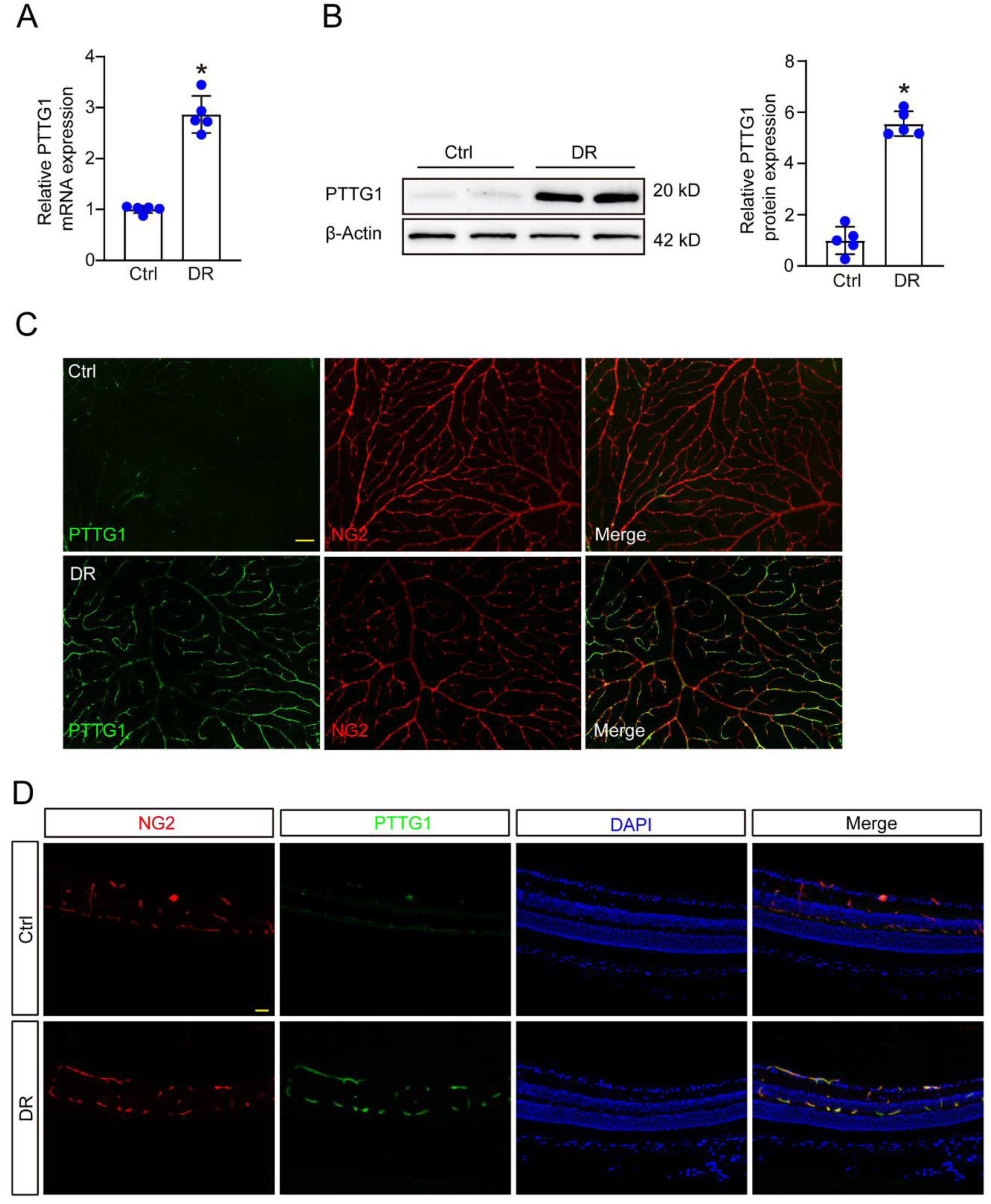
The expression of PTTG1 is upregulated in the retina of DR and co-localizes with pericytes (A and B) qRT-PCR assays and western blot assays were conducted to compare the expression of PTTG1 gene in the retinas of DR mice and non-diabetic mice 6 months after modelling. n = 4; \**P* < 0.05; Student’s *t* test. (C) Immunofluorescence staining assays of retinal flat mounts were performed to assess PTTG1 expression in DR compared with non- DR retinas 6 months after modelling. NG2 was used to label retinal pericytes. Scale bar: 50 µm. (D) Immunofluorescence staining assays of frozen sections illustrate the co- localization of PTTG1 and NG2 in DR and non-DR retinas. DAPI staining was used to label the cell nucleus. Scale bar: 50 µm.

### PTTG1 is involved in high glucose (HG) induced pro-angiogenesis, pericyte mitochondrial dysfunction and apoptosis

Subsequently, we investigated the effects of PTTG1 knockdown on the function of pericytes *in vitro*. Initially, we designed and selected PTTG1 siRNAs with optimal efficiency to knock down PTTG1 expression in pericytes. As western blotting analyses and qRT-PCR showed, the expression level of PTTG1 in transfected cells was effectively knocked down (Figure 4A, B). We exposed pericytes to 50 nm/ml high glucose medium conditions to observe the functional changes after PTTG1 knockdown. In early stages of DR, reduced pericyte recruitment ability is a key factor in the process of pathological angiogenesis.^6–8^ In matrigel co-culture assays, pericytes transfected with PTTG1 siRNA displayed increased recruitment by ECs compared with the Scr siRNA transfected cells in high glucose environments (Figure 4C). During the development of DR, chronic high-sugar environments will promote pericyte apoptosis, which in turn reduces the pericyte coverage area and causes the vessel wall to become more fragile.^23,24^ Calcein-AM/PI staining assays indicated that PTTG1 knockdown attenuated HG-induced pericyte apoptosis, as evidenced by the decrease in PI-positive cells (Figure 4D). During early stages of apoptosis, abnormal changes in mitochondrial structure and function result in the inability to maintain the inner membrane’s transmembrane potential, followed by a decreased potential difference. To detect changes in pericyte mitochondrial membrane potential (MMP), we performed Rhodamine 123 staining and JC-1 staining assays. Both assays demonstrated that, relative to the Scr siRNA group, PTTG1 knockdown alleviated the high-glucose-stimulated MMP collapse in pericytes, allowing the polarization state of MMP to be maintained (Figure 4E, F). We subsequently knocked out the PTTG1 gene in pericytes using CRISPR/Cas9- mediated genome editing (Figure S3A-B). We then repeated the *in vitro* assays previously described, and the results were consistent with those observed following PTTG1 knockdown. (Figure S3C-F). All these results suggested that PTTG1 may play a crucial role in the biological regulation of pericytes *in vitro*.

**Figure 4:**
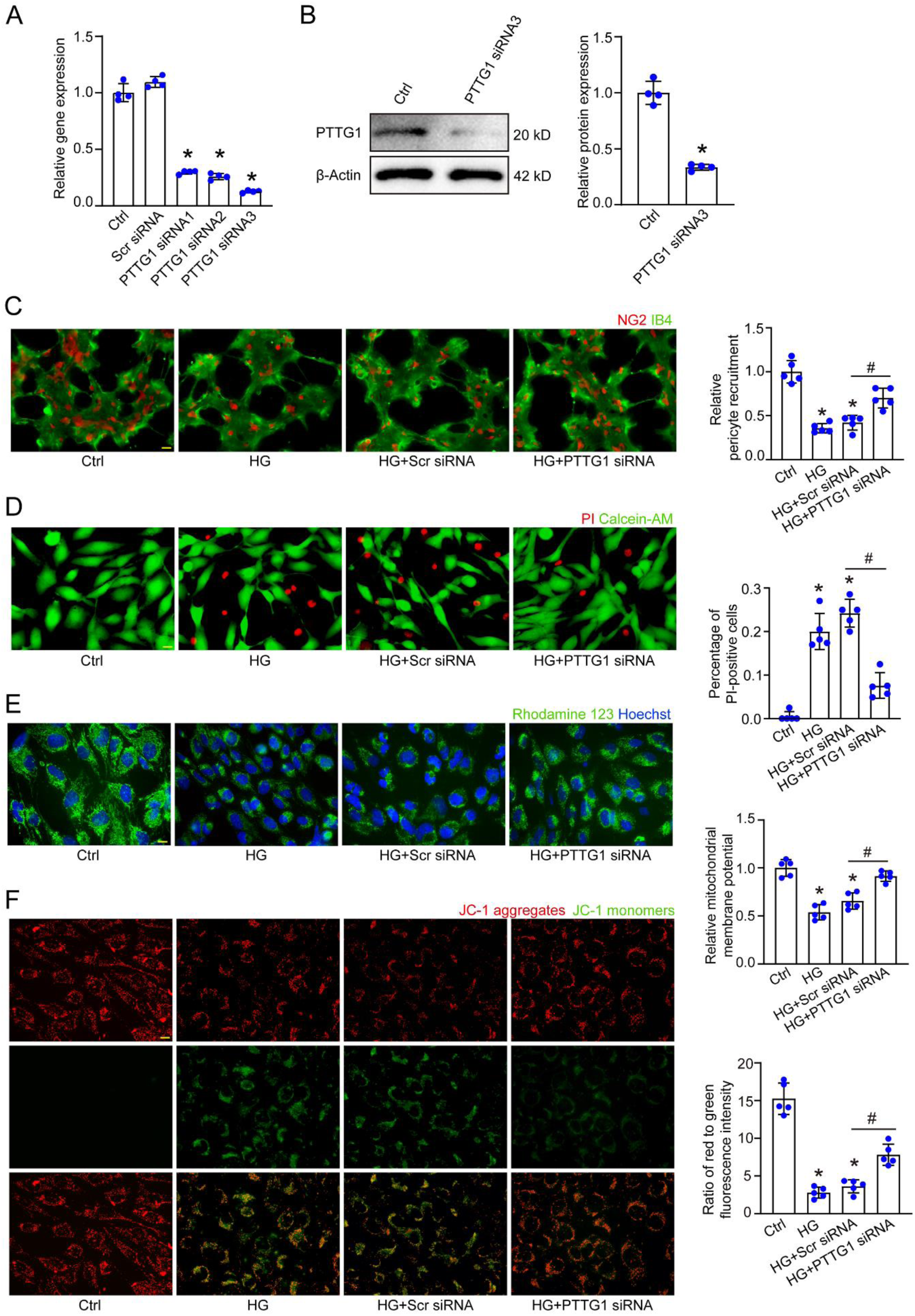
PTTG1 is involved in pericyte biology *in vitro* (A) Pericytes were transfected with PTTG1 siRNA1-3, scrambled (Scr) siRNA, or left untreated (Ctrl) for 24 h, and PTTG1 expression was measured by qRT-PCR. n = 4; \**P* < 0.05; One-way ANOVA with Bonferroni test. (B) Pericytes were transfected with Scr siRNA or PTTG1 siRNA3 for 48 h, and PTTG1 expression was detected by western blotting. n = 5; \**P* < 0.05; Student’s t test. (C - F) Pericytes were transfected with Scr siRNA, PTTG1 siRNA3, or left untreated for 6 h, then exposed to 50 nm/ml high glucose for 48 h. The non-high-glucose group served as the control (Ctrl). Pericyte recruitment to ECs (C) was assessed by NG2 (pericytes) and IB4 (ECs) staining (n = 5, scale bar: 50 µm). Calcein-AM/PI (D) staining detected live (green) and apoptotic (red) pericytes. Mitochondrial membrane potential changes were analyzed using Rhodamine 123 (E) and JC-1 (F) staining assays. Rhodamine 123 (green) and JC-1 aggregates (red), JC-1 monomers (green), Hoechst (blue). n = 5; scale bar: 20 µm; \**P* < 0.05 versus Ctrl; #*P* < 0.05 Scr siRNA versus PTTG1 siRNA. One-way ANOVA with Bonferroni test.

### PTTG1 is a regulator of pericyte shedding and vascular integrity

To investigate the role of PTTG1 in vivo, we established an STZ-induced DR model to examine its effect on retinal vascular dysfunction. We designed three short hairpin RNAs (shRNAs) targeting PTTG1, packaged them into lentiviral vectors, and injected them intravitreally. Seven days later, qRT-PCR and western blotting confirmed the knockdown efficiency, with PTTG1 shRNA3 being the most effective and selected for further experiments (Figure 5A, B). Chronic hyperglycemia causes altered blood flow and increased permeability of retinal vessels with loss or detachment of ECs and pericytes.^5,9,25^ Vascular leakage and decellularized capillary formation are significant features of vascular dysfunction in DR. Evans Blue assays showed that PTTG1 shRNA significantly reduced vascular leakage in diabetic retinas (Figure 5C). Retinal trypsin digestion revealed decreased acellular capillary formation after PTTG1 knockdown (Figure 5D). Immunofluorescent staining of IB4 with NG2 demonstrated that PTTG1 knockdown rescued pericyte loss compared to Scr shRNA-injected mice (Figure 5E).

**Figure 5:**
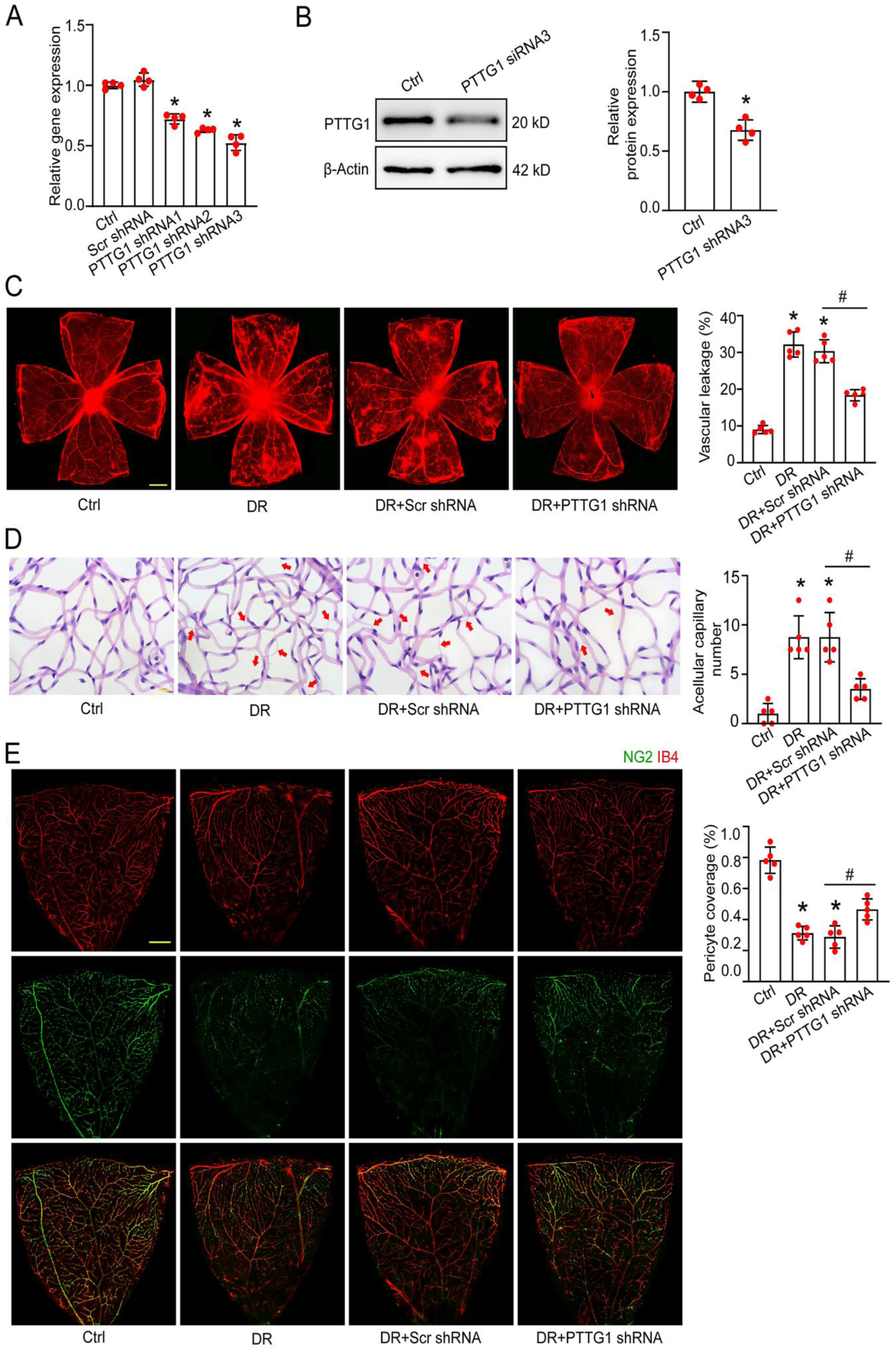
Inhibition of PTTG1 alleviates diabetes-induced retinal vascular dysfunction *in vivo* (A) In STZ-induced diabetic retinopathy (DR) C57BL/6J mice, intravitreal injections of PTTG1 shRNA1-3, negative control (Scr shRNA), or no treatment (DR) were administered for 7 days. The non-diabetic C57BL/6J mice were used as the control (Ctrl) group. PTTG1 expression levels were measured by qRT-PCR. n = 4; \**P* < 0.05; One-way ANOVA with Bonferroni test. (B) For 7 days, DR mice received intravitreal injections of PTTG1 shRNA3 or Scr shRNA. PTTG1 expression was detected by western blot. n = 5; Student’s t-test. (C) Evans Blue assay was performed to assess retinal vascular leakage in 6-month-old DR mice treated with either Scr shRNA, PTTG1 shRNA, or left untreated (DR) for 7 days. Non-diabetic mice were used as the Ctrl group. Evans Blue injections followed. n = 5; scale bar: 500 µm; One-way ANOVA with Bonferroni test. (D) Retinal trypsin digestion was used to detect acellular capillaries, marked by red arrows. n = 5; scale bar: 10 µm; One-way ANOVA with Bonferroni test. (E) Pericyte coverage in the retina was assessed by IB4 and NG2 staining, with NG2 showing green fluorescence and IB4 showing red fluorescence. Scale bar: 100 µm. n = 5; \**P* < 0.05 versus Ctrl group; #*P* < 0.05 Scr shRNA versus PTTG1 shRNA. One-way ANOVA with Bonferroni test.

Single-cell transcriptome sequencing revealed a correlation between upregulated PTTG1 in pericyte subclusters and DR. We designed an adeno-associated virus (AAV) carrying a plasmid to knock down PTTG1 specifically in pericytes. Targeted silencing was achieved by intravitreal injection of AAV into mice. To evaluate cell-specific targeting and silencing efficiency, immunofluorescence staining was performed on retinal sections 21 days post-injection. In Ctrl mice, the green fluorescent protein (GFP) signal from the AAV vector colocalized with the pericyte marker NG2 (Figure S4A). In DR mice treated with PTTG1 AAV, reduced PTTG1 fluorescence intensity confirmed specific downregulation of PTTG1 in retinal pericytes (Figure S4B). Consistent with prior trends, PTTG1 knockdown by AAV mitigated pericyte loss and vascular dysfunction in DR (Figure S4C-E). Given these findings, it is reasonable to assume that the PTTG1 gene plays a pivotal role in diabetes-induced retinal vascular dysfunction and pericyte detachment in vivo.

### Differential analysis of the transcriptome and metabolome of pericytes after knockdown of PTTG1

To identify the alterations in both gene expression and metabolic pathways that occur after knockdown of PTTG1 in pericytes, we performed transcriptomic and GC-MS-based untargeted metabolomic analyses on pericytes transfected with PTTG1 siRNA versus Scr siRNA (Figure 6A). We constructed principal component analysis (PCA) models to investigate the different clustering of different transcriptional profiles. 3D PCA modelling demonstrated the distribution and segregation trends observed after knockdown of PTTG1. As shown, the 3 major components exhibited significant clustering within the 2 groups (Figure 6B). A total of 974 differentially expressed genes were identified by comparing gene expression levels (Fragments per kilobase of transcript per million mapped reads (FPKM > 1.0 and P-value < 0.05). The 525 genes were up-regulated and 449 genes were down-regulated with respect to the Scr siRNA group (Figure 6C). GO terms showed that the top 10 significantly altered terms are mainly involved in cell division, apoptotic process, and regulation of angiogenesis (Figure 6D). Pathway analyses also showed that the knockdown of PTTG1 altered the angiogenesis pathways, the apoptosis-related pathways and the glycolysis-related pathways (Figure 6E). To summarize, analyses uncovered that pericyte angiogenic function and metabolic pathways were altered after PTTG1 knockdown.

**Figure 6:**
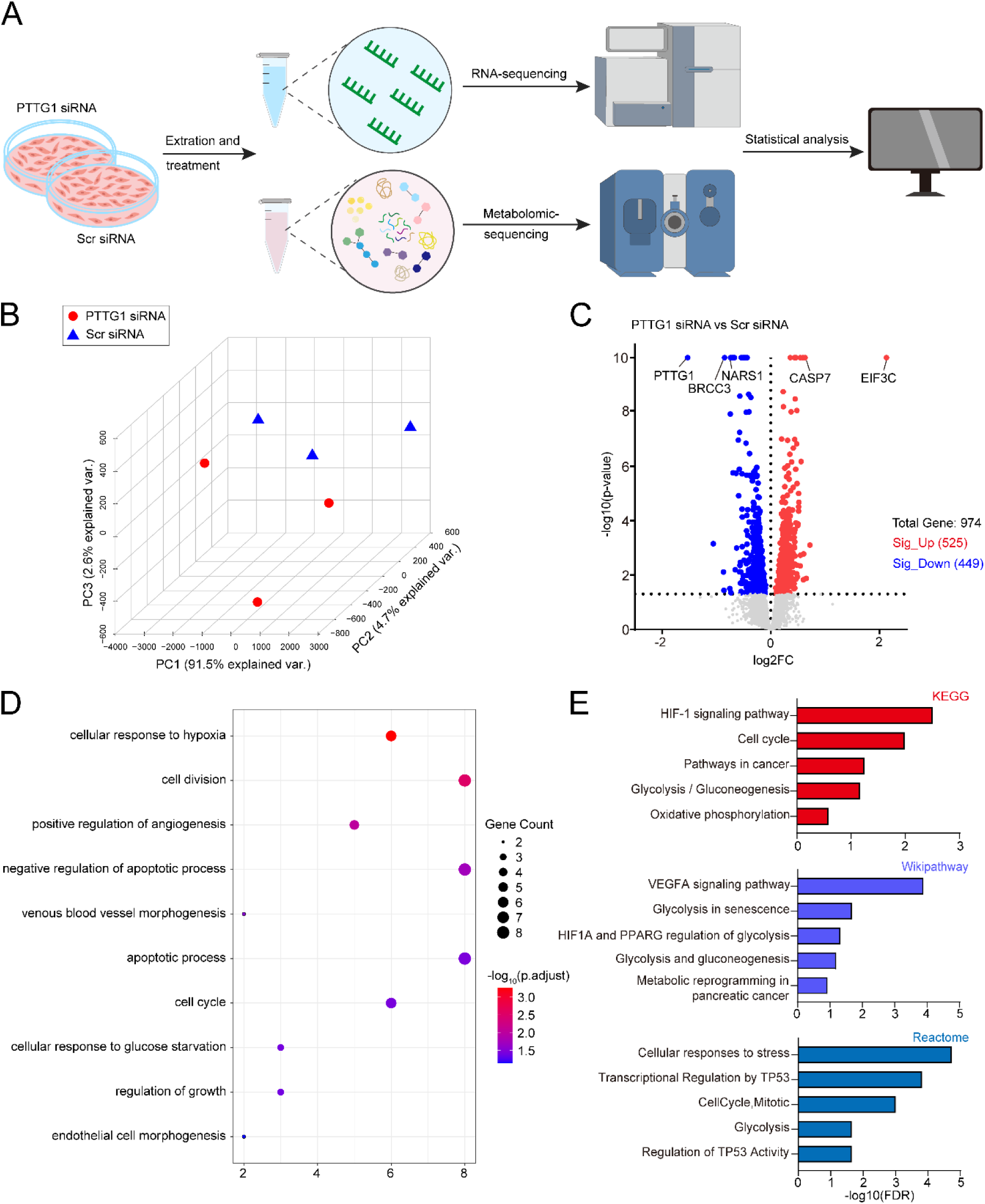
Transcriptomic analysis of pericytes after PTTG1 silencing reveals alterations in gene expression (A) Schematic showing transcriptomic and metabolomic analyses of pericytes transfected with PTTG1 siRNA or Scr siRNA. (B) PCA based on RNA-seq data from pericytes transfected with PTTG1 siRNA and Scr siRNA (n = 3), highlighting PC1, PC2, and PC3 dimensions, with each dot representing a replicate. (C) Volcano plot displaying 525 up- regulated and 449 down-regulated genes between Scr siRNA and PTTG1 siRNA groups, with key genes labelled. (D) GO-BP enrichment analysis of DEGs between PTTG1 siRNA and Scr siRNA groups. (E) Pathway analysis (KEGG, Wikipathway, Reactome) of DEGs between PTTG1 siRNA group and Scr siRNA group..

The differentially expressed metabolites following PTTG1 knockdown were analyzed by mass spectrometry. Orthogonal Partial Least Square Discriminant Analysis (OPLS-DA) identified metabolite differences across experimental groups and generated score plots (Figure 7A). A total of 419 significantly differentially expressed metabolites were detected, with 206 up-regulated and 213 down-regulated (VIP > 1.0, *P*-value < 0.05). Serine and 3- phosphoglycerate were significantly elevated after PTTG1 knockdown (Figure 7B). To accurately elucidate the underlying mechanisms of transcriptomic and metabolic alterations, pathway enrichment analyses of differential metabolites and DEGs revealed that glycolysis and gluconeogenesis were the most significantly altered pathways (Figure 7C, D). The differential gene expression in glycolysis was generally reduced following PTTG1 knockdown (Figure 7E). The pathway diagram highlights genes and metabolites up- or down-regulated in glycolysis and gluconeogenesis (Figure 7F).

**Figure 7:**
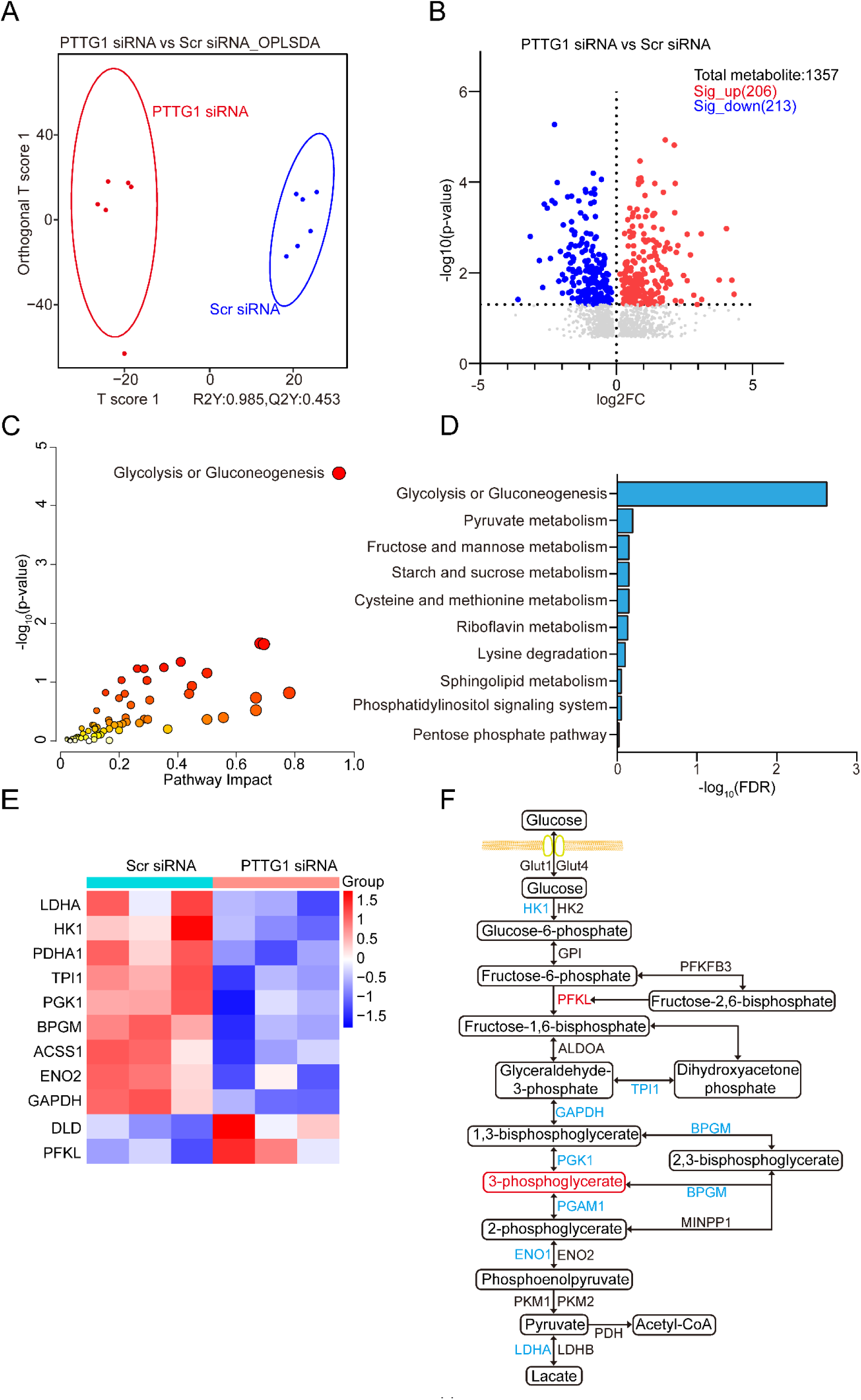
Joint metabolomic and transcriptomic analyses reveal altered glycolytic metabolism in pericytes following PTTG1 silencing. (A) Orthogonal Partial Least Squares Discriminant Analysis (OPLS-DA) was used to assess metabolic differences in pericytes transfected with PTTG1 siRNA and Scr siRNA (n = 6). (B) The volcano plot shows metabolic variations between the PTTG1 siRNA and Scr siRNA groups, revealing 206 up-regulated metabolites and 213 down-regulated metabolites. (C and D) The Joint Pathway Analysis module in MetaboAnalyst 6.0 performed a multi-omics analysis by integrating transcriptome and metabolome data. The volcano plot (C) highlights significantly differential metabolic pathways, with the top 10 enriched pathways listed (D). Glycolysis and gluconeogenesis emerged as the most significantly altered metabolic pathway. (E) The heatmap displays differentially expressed genes (DEGs) in glycolysis and gluconeogenesis pathways between the PTTG1 siRNA and Scr siRNA groups. (F) The pathway diagram shows the upregulation (red) and downregulation (blue) of genes and metabolites involved in glycolytic metabolism after PTTG1 gene knockdown, with metabolites highlighted in boxes.

### Knockdown of PTTG1 in pericytes attenuates glycolytic metabolism and restores mitochondrial function

Prolonged high glucose exposure elevates glycolysis in pericytes, directing intermediates to the polyol and hexosamine pathways and increasing oxidative stress, leading to ROS accumulation, antioxidant depletion, and mitochondrial fragmentation.^10^ To assess PTTG1’s role in glucose metabolism, we measured key glycolytic enzyme expression via western blotting and qRT-PCR (Figure S5A, B). Consistent with enrichment analyses, PTTG1 siRNA reduced HK1, ENO2, and LDHA expression, indicating diminished the pericytes’ ability to catabolize glucose to generate energy with glycolysis. Seahorse assays revealed decreased glycolysis rate, maximum ECAR, and glycolytic reserve capacity following PTTG1 knockdown (Figure S5C-F). Lactate and pyruvate levels, key glycolytic metabolites, were reduced, confirming indeed impeded of glycolysis activity (Figure S5G, H).

High glucose leads to cellular mitochondrial oxidative stress damage, which in turn leads to energy crisis, mitochondria-dependent apoptosis, and ultimately pericyte loss.^9,26,27^ Previous assays on apoptosis and MMP confirmed that knockdown of PTTG1 alleviated the alteration of mitochondrial inner membrane potential difference, so we further explored whether knockdown of PTTG1 could reduce mitochondrial oxidative stress damage and restore biosynthesis. Glutathione is an essential intracellular reducing agent that enhances cellular resistance to reactive oxygen species.^28,29^ High glucose concentrations were observed to reduce intracellular concentrations of glutathione (GSH) and nicotinamide adenine dinucleotide (NADH). In contrast, knockdown of PTTG1 was found to elevate levels of GSH and NADH, thus increasing cellular antioxidant capacity. Using the reactive oxygen species fluorescent probe DCFH-DA to detect intracellular ROS, we found that knockdown of PTTG1 alleviated the elevated ROS levels from high glucose (Figure S6A- F). Subsequently, Seahorse assay was employed for oxygen consumption rate (OCR) to investigate alterations in mitochondrial synthetic functionality. The results demonstrated that, after PTTG1 knockdown, pericyte basal respiration remained unaltered, yet maximal respiration was enhanced, accompanied by increased ATP production by mitochondria. Proton leakage from the cells was also reduced, indicating recovery from mitochondrial damage (Figure S6G-K). Based on these results, it is evident that PTTG1 downregulation in pericytes alleviates high glucose-stimulated mitochondrial oxidative stress and protects mitochondrial function.

### In clinical proliferative retinal diseases, PTTG1 is highly expressed in neovascular and pericyte locations

To investigate PTTG1 dysregulation in retinal vascular diseases, we analyzed clinical samples from proliferative diabetic retinopathy (PDR) and epiretinal membrane (ERM) patients. In PDR, fibrovascular membranes form during pathological neovascularization, leading to vitreous hemorrhage or retinal detachment.^5,30^ Immunofluorescence revealed PTTG1 expression in PDR fibrovascular membranes, co-localizing with the pericyte marker NG2, but not significantly expressed in ERM (Figure 8A, B). In retinopathy of prematurity (ROP), where disorganized vasodilatation causes hemorrhage and scarring^20^, PTTG1 expression was elevated in retinal neovascularization areas of OIR mice and co- localized with NG2 in ROP proliferative membranes (Figure 8C, D). These findings indicate that PTTG1 dysregulation contributes to the pathogenesis of retinal vascular diseases.

**Figure 8.**
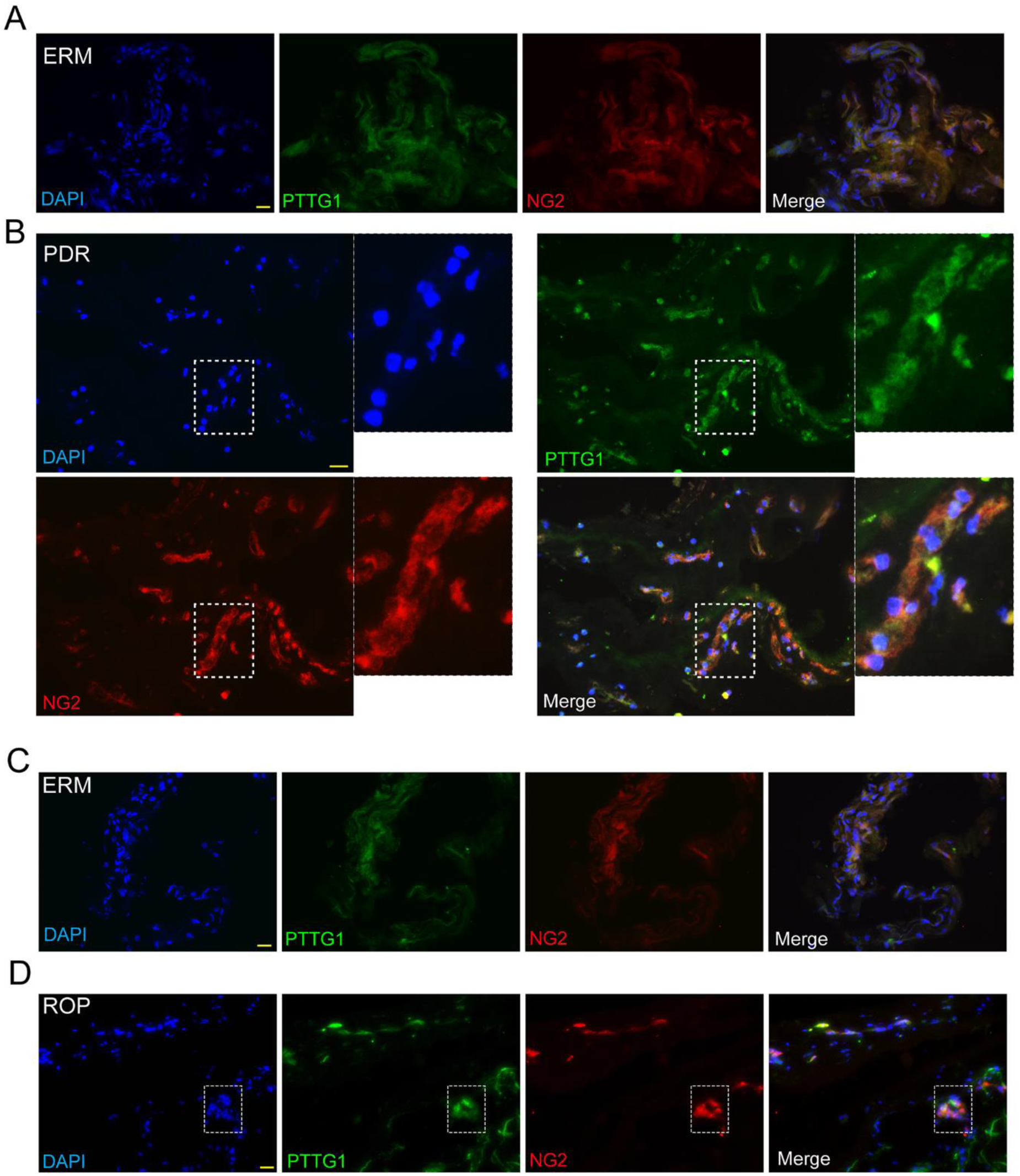
Clinical correlation between PTTG1 and retinal neovascularization diseases (A-D) Immunofluorescence assay to detect PTTG1 expression in fibrous membranes of patients with ERM, and fibrovascular membranes of patients with PDR or ROP. Scale bar: 50 µm. PDR, proliferative diabetic retinopathy. ROP, retinopathy of prematurity. ERM, epiretinal membrane.

## Discussion

DR is a disease characterized by retinal microvascular dysfunction and degeneration. Vision loss is often accompanied by early-stage retinal edema, hemorrhages, exudates, and microaneurysms, progressing to neovascularization and fibrovascular membrane proliferation in later stages.^31,32^ Research on DR treatment primarily focuses on VEGF-based therapies and the management of complications following neovascularization in advanced stages. However, treatment strategies targeting early-stage microvascular damage are frequently overlooked.^33,34^ Focal pericyte degeneration and loss, as initiating events in many microvascular pathologies, have been recognized and further investigated in fields such as diabetic nephropathy,^35^ hypertensive encephalopathy,^36^ and chronic obstructive pulmonary disease.^37^ However, the mechanisms of pericyte dysfunction in studies of DR are still not fully understood. In this study, we identified a subcluster of pericytes in DR retinas characterized by the high expression of the PTTG1 gene through single-cell RNA sequencing. These pericytes were closely associated with retinal vascular formation. Silencing PTTG1 preserved pericyte function and delayed retinal vascular dysfunction, indicating that this gene is a potential target for treating microvascular complications in DR.

Pericytes, termed “mural cells” upon discovery, encase endothelial cells and form tight connections via “peg-and-socket” junctions or adherens junctions, where microfilaments on the plasma membrane embed into adjacent endothelial cell cytoplasm.^38^ These connections allow pericytes to regulate the microvascular system, supplying nutrients and oxygen to retinal cells and supporting visual function.^39^ Diseases and stress conditions like hypoxia, inflammation, and hyperglycemia can trigger pericyte migration, degeneration, disrupted cell-cell communication, endothelial apoptosis, and pathological proliferation, damaging the retinal microvasculature and exacerbating disease progression.^7,8^ Pericytes exhibit notable inter- and intra-tissue heterogeneity due to their dual origin from neuroectoderm and mesoderm, resulting in tissue-specific subclusters.^40^ In the CNS, pericytes densely encase endothelial cells, forming the BBB, with NG2 highly expressed during development and CD13 in the mature BBB. The retina, as an extension of the CNS, mirrors this distribution, while cardiac and skeletal muscle pericytes resemble smooth muscle cells, expressing α-SMA.^41^ Hepatic pericytes (stellate cells) associate loosely with sinusoidal endothelium, functioning in Vitamin A storage.^42^ Comprehensively understanding pericyte heterogeneity is crucial for providing new insights into the mechanisms of vascular dysfunction and the targeting of pathological vasculature for therapy. Advances in single-cell RNA sequencing have unveiled pericyte subpopulations. Muhl *et al.*^43^ identified fibroblast and pericyte heterogeneity across organs, while Li *et al* ^44^ discovered a novel TCF21-expressing pericyte promoting colorectal cancer metastasis. However, Studies investigating how the abundant pericytes in the retina regulate vascular formation through differential gene expression from a single-cell perspective are limited.

Our research employed single-cell RNA sequencing to construct a comprehensive cellular atlas of the retina in DR mice, identifying ten different cell clusters using specific markers. Within the pericyte cluster, we identified three distinct pericyte subpopulations, each with unique gene expression profiles. Subcluster 1 highly expressed key genes such as Tulp1, Hsp90aa1, and Pla2g7, which have been reported to be involved in stimulating phagocytosis,^45^ cellular response to environmental stress,^46^ and regulation of inflammasome activity.^47^ Subcluster 2 showed high expression of genes like Tagln, Acta2, and Tpm2, which are associated with maintaining normal vascular function.^48,49^ Pericyte subcluster 0 specifically expressed angiogenesis-related genes such as Spock2, Cxcl12, Flt1, and Klf4. Enrichment analysis indicated that this subpopulation is closely related to “angiogenesis,” “cell-cell adhesion,” “endothelial tube morphogenesis,” and “positive regulation of cell proliferation.” Therefore, we speculate that pericytes in subcluster 0 may be highly associated with pathological angiogenesis in diabetic retinopathy. Within subcluster 0, PTTG1 was identified as the most significantly upregulated gene in the diabetic group, prompting further investigation into its role.

PTTG1 is an oncogene that plays a crucial role in various biological processes, including cell replication, DNA damage repair, and metabolic regulation. Previous studies have reported its high expression in pituitary, liver, gliomas, and gastrointestinal tumors. PTTG1 is a key gene associated with tumor metastasis and is closely related to endocrine diseases. Zhi *et al.*^50^ confirmed that silencing PTTG1 inhibits the proliferation, migration, and invasion of glioma cells, and this pro-migratory and proliferative function is also observed in aortic vascular smooth muscle cells. Other studies have shown that PTTG1 inhibits the expression of inflammatory factors through the MAPK pathway and protects the extracellular matrix.^51^ In liver fibrosis research, interference with PTTG1 was shown to reduce fibrosis area and decrease α-SMA expression.^52^ This enhanced migratory function and cellular transformation are common in the activity of pericytes in DR. High expression of PTTG1 also stimulates the secretion of fibroblast growth factor (FGF) and VEGF, thereby inducing angiogenesis.^53,54^ Our study demonstrates that PTTG1 is upregulated in DR and is specifically localized to pericytes. In the OIR model, PTTG1 is highly expressed in neovascular tufts. Silencing PTTG1 rescued pericytes from reduced recruitment by endothelial cells, decreased mitochondrial membrane potential, and increased apoptosis under high glucose conditions. Knockdown of PTTG1 in mouse retinas alleviated pathological vascular leakage and the formation of acellular capillaries, partially restoring pericyte coverage. These findings indicate that PTTG1 regulates pericyte function in the DR retina.

In pancreatic cancer, PTTG1 overexpression promotes cell proliferation and the Warburg effect by regulating the c-myc pathway.^55^ Glycolysis plays a critical role in pericyte function across organs. Meng et al.^56^ showed that increased glycolysis via hexokinase 2 (HK2) in tumor pericytes enhances ROCK2-MLC2-mediated contractility, impairing vascular support. Reducing glycolysis by targeting the HIF1α-HK2 pathway or lowering pyruvate kinase M2 (PKM2) expression can inhibit the pericyte-myofibroblast transition, slowing acute kidney injury progression.^10,57^ Our transcriptomics and metabolomics analysis revealed that silencing PTTG1 downregulated glycolysis-related genes, reducing energy production, cell activity, and glycolytic metabolites. The general downregulation of related genes reduced energy production via glycolysis, leading to decreased cell activity and fewer glycolytic metabolites. Long-term high glucose environments have been reported to increase glycolysis levels, redirecting intermediates to the polyol and hexosamine biosynthesis pathways, enhancing NAD(P)H oxidase activity, and increasing cellular oxidative stress. Reduced intracellular reductants deplete antioxidant defenses, enhance protein glycation, and cause mitochondrial fragmentation.^58^ Our study shows that PTTG1 knockdown decreased ECAR and reduced glycolytic metabolites lactate and pyruvate, while intracellular reductants NADH and GSH levels increased, suggesting enhanced resistance to high glucose-induced oxidative stress. Intracellular ROS levels decreased, and mitochondrial morphology and membrane potential improved. Interestingly, when PTTG1 was knocked down, the overall OCR levels slightly increased. We do not believe this indicates a metabolic reprogramming from glycolysis to mitochondrial oxidative phosphorylation, as this would further elevate ROS levels, and we did not observe changes in acetyl-CoA or TCA cycle metabolites. Some studies suggest that PTTG1 knockdown may lead to a concurrent decrease in glycolysis and mitochondrial synthesis.^55^ We hypothesize that, in the absence of cellular stress, this is a slight energy compensation by the mitochondria for the reduction in glycolysis, with restored mitochondrial function and reduced proton leakage in the proton transport chain. Research on the metabolic mechanisms of pericytes in the retina is a crucial direction for the future, potentially addressing the issues of frequent injections and resistance associated with anti-VEGF drugs.

In this study, we characterized the heterogeneity of pericytes in the DR process through single-cell RNA sequencing. We identified a pericyte subpopulation with high expression of the PTTG1 gene, which is closely associated with angiogenic functions and shows significantly elevated expression in DR tissues. Silencing PTTG1 can rescue pericyte dysfunction *in vitro*, reduce apoptosis, and alleviate pathological angiogenesis of DR *in vivo*. Silencing PTTG1 also downregulates the expression of genes related to the glycolysis pathway, reducing oxidative stress levels in pericytes under high-glucose conditions. PTTG1 can serve as a metabolic target for abnormal angiogenesis, providing insights for the development of new therapies to improve current treatment strategies.

## Sources of Funding

This study was funded by the grants from National Natural Science Foundation of China (no.81770945 and 81970809 to Dr Yan).

## Disclosures

None.

## Supplemental Material

Extended Materials and Methods Figures S1–S6

Tables S1–S2

Major resources table References 59–60

---

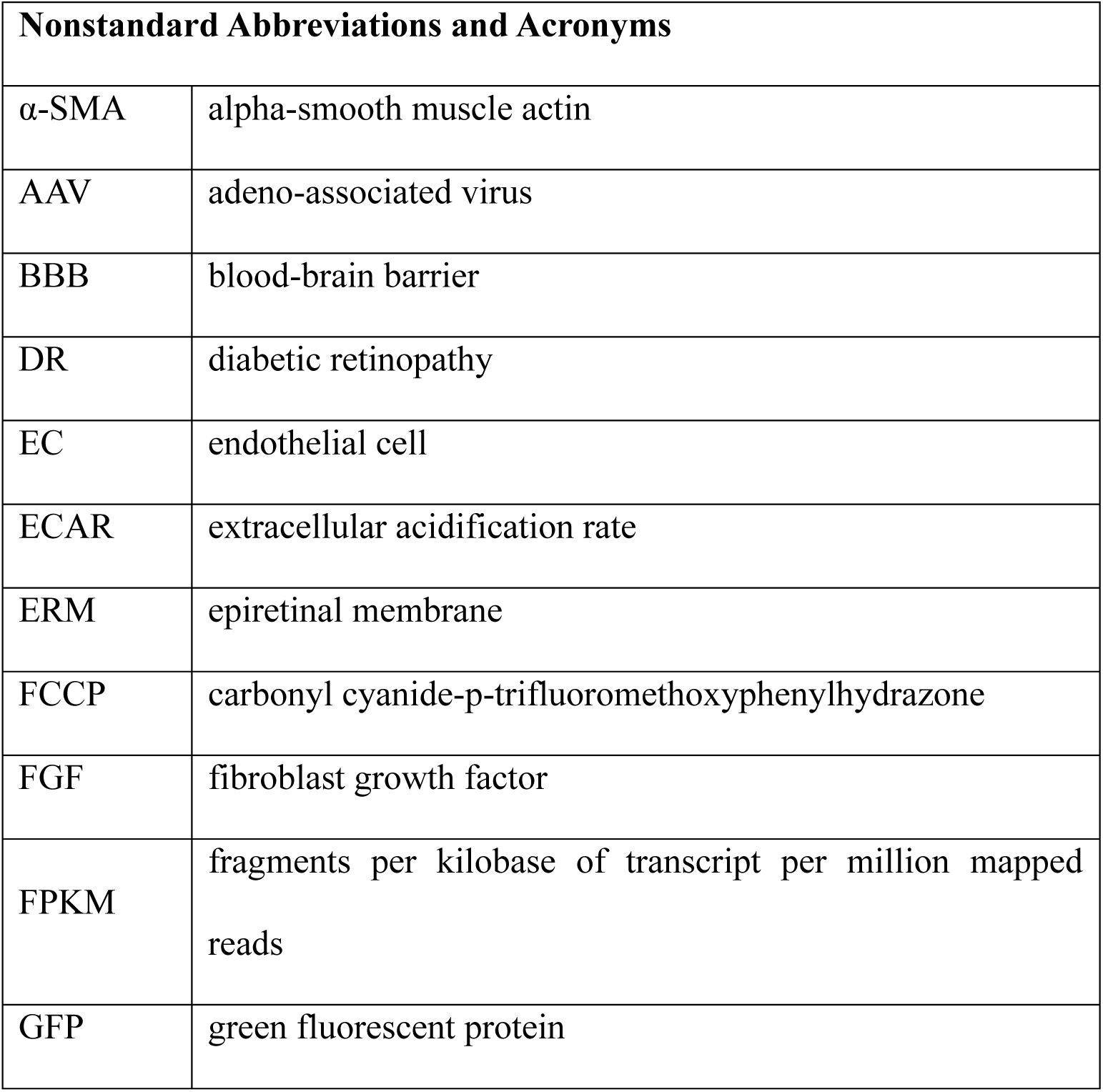
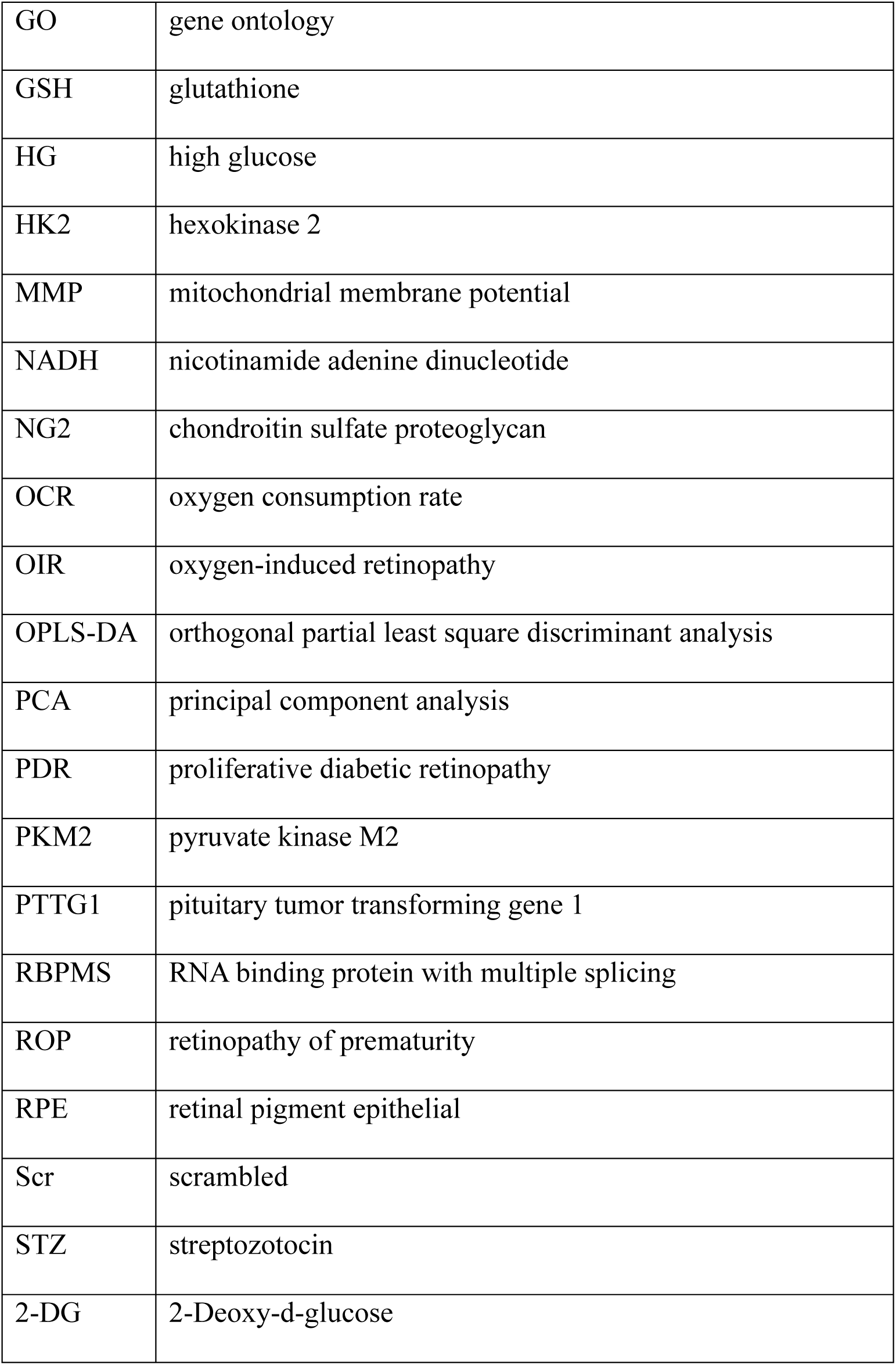

## Novelty and Significance

### What Is Known?

1. Microvascular dysfunction is a key driver of the severe systemic complications associated with diabetes
2. Pericyte loss is one of the earliest events that compromise microvascular integrity in diabetes
3. The functional heterogeneity of pericytes is essential for vascular development and remodeling

### What New Information Does This Article Contribute?

1. Single-cell RNA sequencing of retinal samples identified a novel pericyte subcluster associated with diabetes-induced microvascular dysfunction, characterized by high expression of the PTTG1 gene.
2. PTTG1 promotes pericyte apoptosis, disrupting vascular stability and driving pathological angiogenesis.
3. Silencing PTTG1 restores retinal vascular function by modulating glycolytic metabolism in pericytes

Diabetic retinopathy (DR) is a major microvascular complication of diabetes, with pericytes playing a critical role in maintaining vascular integrity. This study examines the transcriptomic diversity of retinal pericytes using single-cell RNA sequencing, identifying three distinct pericyte subclusters. Subcluster 0, which is associated with microvascular complications, exhibits elevated expression of pituitary tumor-transforming gene 1 (PTTG1). Mechanistic investigations show that diabetic stress induces PTTG1 expression in pericytes, contributing to vascular dysfunction through metabolic reprogramming and increased apoptosis. Silencing PTTG1 mitigates these effects, positioning it as a promising therapeutic target. These findings expand our understanding of pericyte biology in DR and suggest that targeting specific pericyte subclusters and their associated metabolic pathways could provide novel strategies for preventing or treating diabetic microvascular complications.

## Notes

### Competing Interest Statement

The authors have declared no competing interest.

